# A flexible, efficient, and scalable platform to produce circular RNAs as new therapeutics

**DOI:** 10.1101/2022.05.31.494115

**Authors:** Huanhuan Wei, Chuyun Chen, Kai Zhang, Zeyang Li, Tong Wei, Chenxiang Tang, Yun Yang, Zefeng Wang

## Abstract

Messenger RNA (mRNA) has recently emerged as a new drug modality with great therapeutic potential. However, linear mRNAs are relatively unstable and require base modification to reduce their immunogenicity, imposing a limitation to the broad application. The circular RNA (circRNA) presents a better alternative for prolonged expression of the proteins. However the *in vitro* circularization of RNA at industrial scale is technically challenging, and the direct comparison of efficacy between the circRNA and linear mRNA drugs is lacking. Here we developed a new self-catalyzed system to efficiently produce circRNAs in a co-transcriptional fashion. By rational sequence design, we can efficiently produce scarless circRNAs that do not contain foreign sequences. The resulting circRNAs are very stable and have low immunogenicity, enabling prolonged protein translation in different cells without cellular toxicity. The circRNAs encapsulated in lipid nanoparticles can be efficiently delivered into mice to direct robust protein expression with improved duration and efficiency. Finally, the circRNAs encoding RBD of SARS-CoV-2 S protein induced strong antibody productions, with neutralization antibody titers higher than the preclinical data from the linear mRNAs. Collectively, this study provided a general platform for efficient production of circRNAs with robust activity in directing protein production, demonstrating the potential of circRNAs as the new generation of mRNA therapy.

## Introduction

mRNA has recently come into focus as a new drug modality with great therapeutic potential, as mRNA-based therapy can theoretically use any protein as the active pharmaceutical ingredient ^1^. Despite the clear demonstration of efficacy for infectious disease and cancer vaccines, application of mRNA for non-vaccine therapeutics has been limited by the duration of expression, stability, immunogenicity, and the ability to control cell type-specific expression. In addition, there are technical challenges associated with mRNA production, modification, and extra-hepatic delivery. Circular RNAs (circRNAs) are covalently closed RNA molecules generated mainly from pre-mRNA back-splicing ^2^, and have also been found to function as mRNAs to direct protein synthesis inside cells ^3^. As a new generation of therapeutic mRNAs, circRNAs may address some of the limitations of linear mRNA including increased stability and reduced immunogenicity ^4,5^. Therefore, translatable circRNAs have the potential to expand the application of mRNA-based therapy, especially when a prolonged expression of the target protein is required. A major limitation for the broad application of circRNAs is the *in vitro* production of circRNAs, especially in the step of RNA circularization after the *in vitro* transcription (IVT)^6^. The most commonly used methods for RNA circularization include the end-ligation of linear RNA with T4 RNA ligase ^7^, or the permuted intron-exon (PIE) splicing strategy using the self-catalyzed group I introns ^8,9^. However the direct ligation with RNA ligases usually needs to use a “splint” sequence to connect the free ends, and thus is technically challenging to scale up the production due to the formation of concatemers ^6^. Moreover, introducing the ligase protein will also increase the complexity in the product purification. On the other hand, the PIE method will introduce a foreign fragment into the circRNA as a “scar” sequence ^9,10^, which may distort the structures of circRNAs to provoke innate immune responses ^11^ and complicate the future approval for therapeutic applications. Recent modifications of the PIE method could produce the “scarless” circRNAs by using the pairing of internal sequences within the circRNAs ^12,13^, thus are less flexible in their sequence choices. Therefore reliable circRNA production methods are needed for the therapeutic application of circRNAs on a large scale. In addition to the technical challenges on circRNA production, the translation of circRNAs is mediated through a cap-independent pathway, which is usually less efficient compared to canonical cap-dependent translation. For the development of successful circRNA drugs, optimization of circRNA translation efficiency is required.

In this study, we developed a new and scalable technology to engineer and produce circRNAs that efficiently direct protein translation. Our design is based on the self-splicing group II intron to achieve co-transcriptional circularization of scarless circRNAs with high efficiency. The resulting circRNAs can be purified and transfected into cells to direct robust cap-independent translation from the internal ribosomal entry sites (IRESs). Using this platform, we generated a series of circRNAs containing different genes to direct the translation in cultured cells and in mice, and the purified circRNAs did not cause deleterious innate immune responses. The therapeutic application of this system was exemplified by engineering circRNAs that encode the RBD of the SARS-CoV-2 S protein to induce robust antibody production in mice. Collectively, this study provided an efficient and scalable platform for circRNA engineering and production, which has broad application in the new generation of mRNA therapy.

## Results

### Design and production of circular RNAs *in vitro*

To efficiently produce circular RNAs, we take advantage of the self-catalyzed splicing reaction by group II introns, which are mobile genetic elements found mainly in bacterial and organellar genomes ^14^. All group II introns have six structural domains (D1 to D6), of which the domain 1 (D1) is the largest domain and contains several short exon binding sites (EBS) to determine the splicing specificity (Fig. 1A). Specifically, the EBS1 in D1 can pair with the intron binding site (IBS1) at the 5’-splicing junction to determine the exact position of splicing, whereas the 3’-splicing junction is determined by the paring of IBS3 with the EBS3 in D1 (for group IIB, IIC and IIE/F introns) or the δ base of D1 (for group IIA introns). The domains 2 and 3 play key roles in assembling the active intron structure and stimulating splicing reaction, whereas the D4 is a stem-loop structure with a long loop containing the ORF of maturase. The highly conserved D5 is the heart of active site for self-splicing ^15^, whereas the D6 contains a bulged adenosine as the branching site (Fig. 1A). The *in vitro* self-splicing of group II intron requires only correct folding of intronic RNA structure and Mg^2+ 16^, however the *in vivo* splicing requires the assistance of the maturase ^17^.

**Fig. 1.**
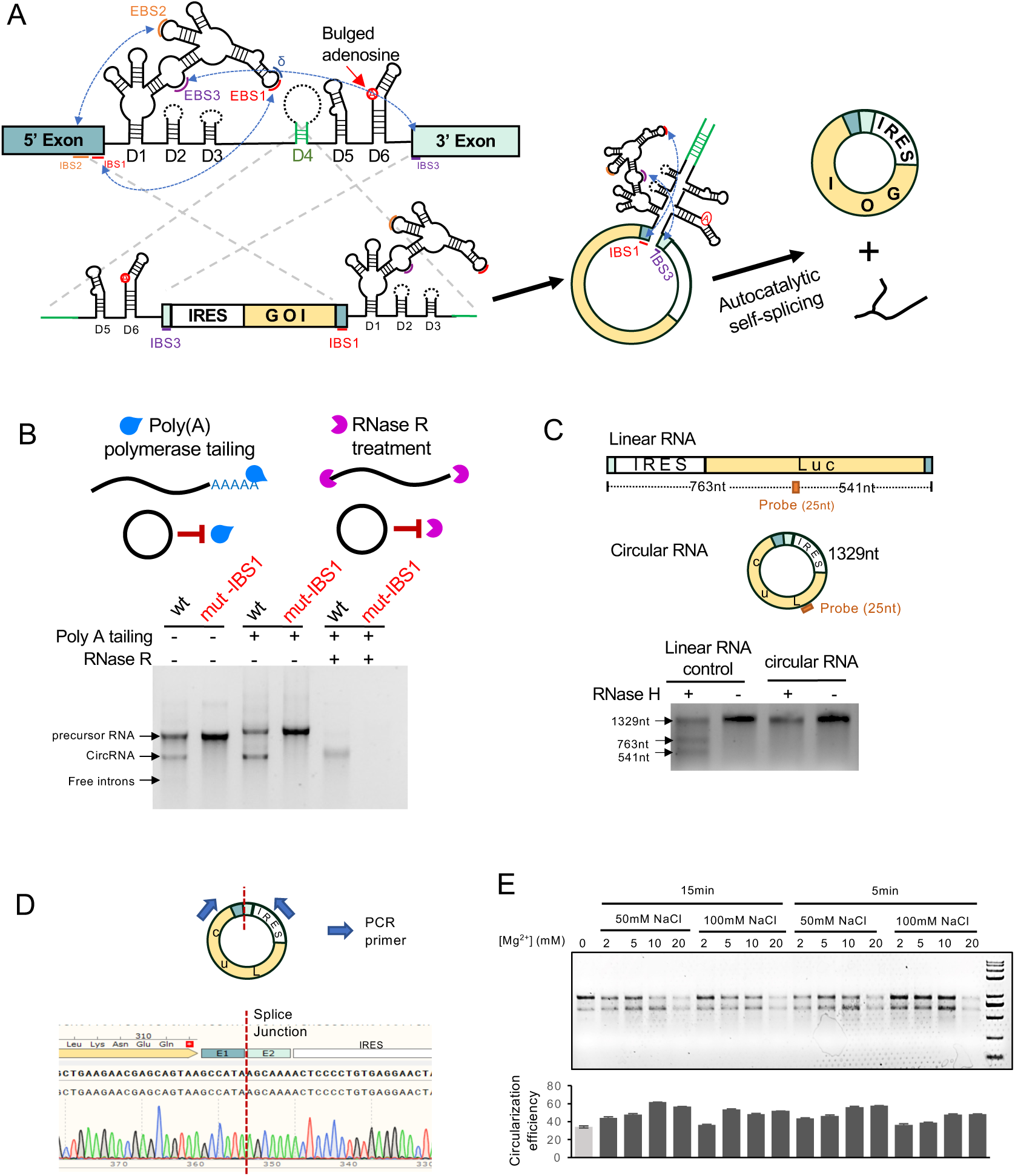
Engineer of a new circRNA production system. (A) Schematic diagram for the design of CirCode system. The autocatalytic self-splicing group II intron was split into two fragments at the D4 domain, and a customized exons containing IRES and coding region of a gene of interest (GOI) were inserted between the split intron. (B) The circRNA is in vitro synthesized and analyzed with agarose gel. IBS1 was mutated to disable self-splicing. After IVT, circularized RNAs were confirmed by Poly A tailing and RNase R treatment. (C) Further confirmation of RNA circularization by RNase H nicking assay. Linear RNA containing the same sequence in circular RNA could be digested into 3 fragments by RNase H. (D) Sanger sequencing output of RT-PCR across the splice junction of the CircRNA sample depicted in lane 1 and lane 3 from (B). (E) Agarose gel demonstrating the effect of cation concentration and reaction time on splicing. Column purified RNA from IVT was incubated with buffer including indicated magnesium (50 mM Tris-HCl, 50 /100 mM NaCl, 0-20 mM MgCl2, pH=7.4) for 5 or 15 min.

Based on this domain configuration, we first split the group II self-splicing intron from the surface layer protein of *Clostridium tetani* (ctSLP) ^18^ at the loop region of different domains (loops of D1, D3, D4), generating a series of split-intron systems that contain a customized exon flanked by the two half-introns (Fig. S1 and Fig. 1A). A fraction of the stem region was separated and placed into each end of the resulting RNA, thus forming a complementary structure that helps the folding of the active intron (Fig. S1 and Fig. 1A, indicated with green lines). Two short 6-nt sequences in the exons, IBS3 (intron binding site 3) and IBS1, were included at each splicing junction to provide long-range interactions with the EBS3 and EBS1 of the intron. Upon the *in vitro* run-off transcription, the resulting RNA precursor could be self-spliced by the ctSLP group II intron to produce a circular RNA of the customized exon and a branched intron RNA. By including an IRES sequence at upstream of a gene of interest, the resulting circular RNA can function as an mRNA to direct protein synthesis through cap-independent translation (Fig 1A). We named this system as the circular coding RNA (CirCode), which could serve as a general platform to produce any given circRNA for protein translation.

To validate this design, we included the ORF of the Renilla luciferase gene together with an IRES into the customized exon, and used IVT to generate this intron-exon-intron RNA precursor. We observed that all the resulting RNA precursors (D1, D3, or D4 split intron) can be self-spliced *in vitro* to produce an extra band corresponding to circRNAs, with the D4 split intron being the most active (Fig. S1). This design was therefore selected for further study. We first mutated the splice junction (at the IBS1) of the D4 split intron, and found that the mutated RNA precursor failed to produce the circRNAs (Fig. 1B). Next, we used two different methods to confirm that the additional band below the precursor RNA is indeed a circRNA, which cannot be extended by tailing reaction by poly-A polymerase and is resistant to the digestion by RNase R treatment (Fig. 1B). In addition, we gel-purified the circRNAs, validated its identity using RNase H digestion (Fig. 1C) and direct sequencing across the junction (Fig. 1D). Finally, we optimized the reaction conditions using different concentrations of MgCl_2_ and NaCl in the circularization buffer following the IVT step (see method), and found that using 10-20 mM Mg^2+^ with 50-100 mM NaCl is the optimal reaction condition for the RNA circularization (Fig, 1E). We also observed certain degree of RNA circularization (∼30%) even when the circularization buffer does not contain any Mg^2+^, presumably because the RNA circularization happened co-transcriptionally in the IVT buffer containing 24mM MgCl_2_ (see methods).

### Platform optimization of circRNA production and translation

The PIE methods using group I introns for circRNA production also introduced an extraneous sequence from T4 bacteriophage or Anabaena into the final products ^9,10^. This “scar” sequence is usually around 80-180 nt long, which limits the design flexibility of target circRNAs and may introduce some unwanted side-effect during drug development. Subsequent methods for “scarless” PIE usually require the pairing of internal sequences within the circRNA product, thus limited the choice of RNA sequences. Our initial design used two short (6-nt) sequences, IBS1 and IBS3, for the intron-exon recognition, which leaves a shorter “scar” of 12-nt. To reduce the potential interference by the scar sequence, we further modified the design by changing the exon binding sites in the D1 domain, making the EBS1 and sequence upstream of EBS1 to respectively form base pairs with the 3’ and 5’ end of the circular exon (Fig. 2A, left). This new design enabled the generation of “scarless circRNAs” without extraneous sequences. We validated this design by generating two different circRNAs encoding the EGFP and Rluc-P2A (Fig. 2A, right), and observed the efficient circularization of scarless circRNA in different ion conditions. The scarless self-splicing of circRNAs was further confirmed by the sequencing of final circRNA products (Fig. 2A, bottom of the right). In summary, we have engineered the CirCode system that can efficiently produce scarless circRNAs of any sequences, providing a general platform for using circRNA as different therapeutic tools. The previous study using the PIE method suggested that addition of a short spacer region before the IRES may assist the correct folding of the IRESs and the active introns ^9^, and thus we introduced several versions of spacer sequences containing IRES-like short elements ^19^ at each end of the circular exon to optimize their circularization and translation efficiency. We found that different spacers indeed affected the RNA circularization efficiency, which ranged from 47% (for SP5) to 83% (for SP4) (Fig. 2B). As expected, different spacers containing short IRES-like elements also affected the translation activities of circRNAs (Fig. S2). We further transfected two circRNAs with different spacers into three cell lines at different doses, and observed dose dependent increase in protein production as judged by the luciferase activity assay (Fig. 2C). Finally, we examined the time course of protein production upon circRNA transfection, and found that the active protein can be detected in six hours and reach the expression peak of expression at 48 hours after transfection (Fig. 2D).

**Fig. 2.**
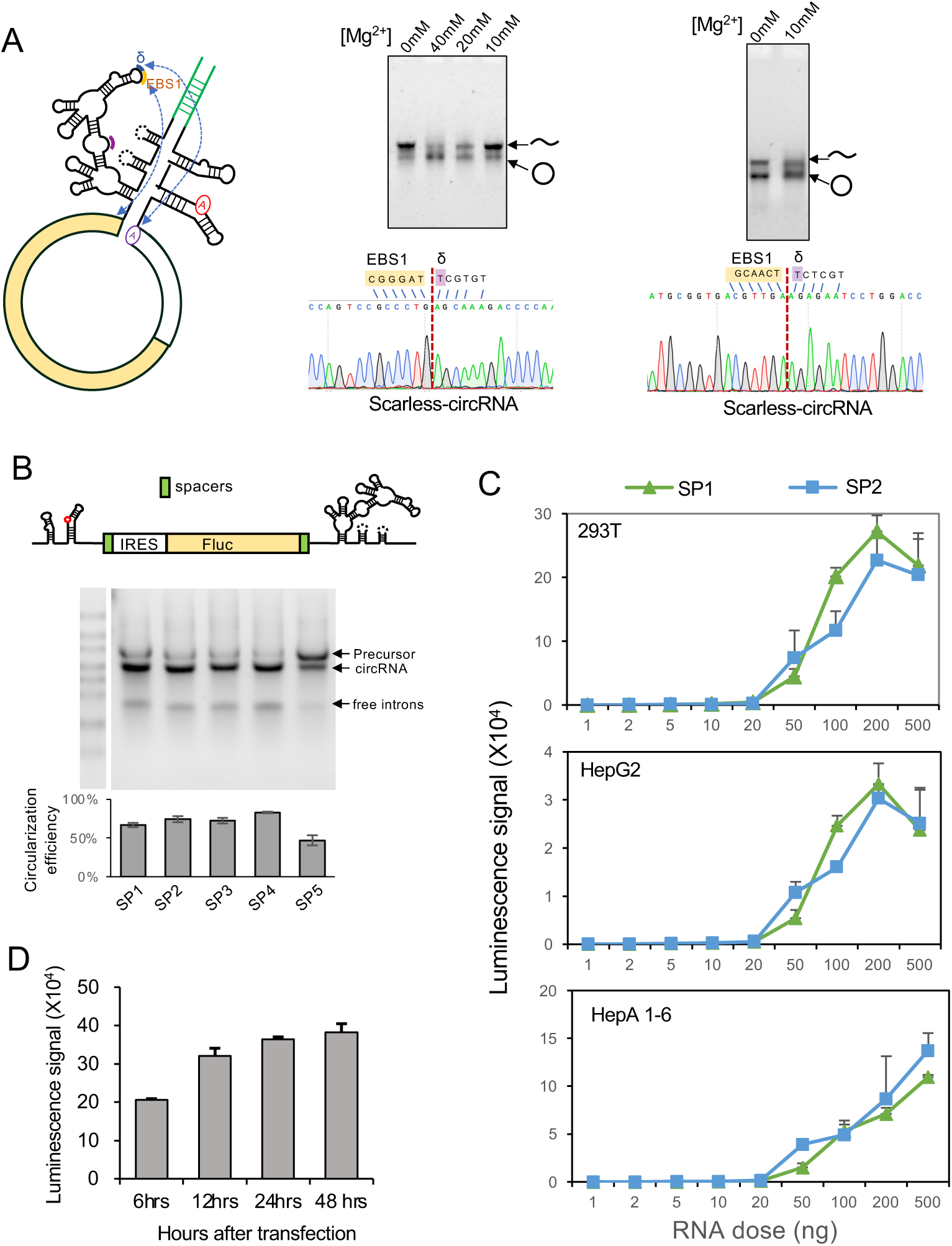
Optimization of circRNA production. (A) Design of scarless circRNA production. Left, the schematic diagram for modification of intronic sequences, resulting in the specific recognition of the splice junction of custom exon and the generation of a scarless circRNA. Right, two examples of scarless circRNAs produced using the CirCode design. The circularization was confirmed by both gel electrophoresis and sequencing of the junction region. (B) Testing different spacer regions between the ORF and the IRES. (C) Transfection and translation of circRNA in different doses. The circRNAs containing two different spacers were gel purified and transfected into three cell lines cultured in 24-well plate at different doses, the activity of firefly luciferase were measured 24 hours after transfection. (D) Time course of circRNA translation. The circRNAs encoding the Rluc gene were transfected into transfected into 293T cells (500 ng circRNAs were used in each transfection), and the luciferase activity were measured at 6, 12 and 24 hours after transfection.

### circRNAs mediate prolonged protein production

A major advantage of circular mRNAs is their superb stability because of lacking the free ends, therefore the circRNA should have a good shelf life for protein expression compared to the linear counterparts. To directly test this, we synthesized both linear and circular mRNA encoding the Gaussia luciferase (Gluc), and stored them parallelly in nuclease-free water at room temperature for different days before transfecting them into 293T cells. We found that the activity of the circRNA to direct protein translation is essentially unchanged during the two weeks, whereas the linear mRNA lost about half of its activity by day 3 of the storage (Fig. 3A), suggesting that circRNAs can be stably stored in room temperature. In addition, we found that once transfected into cells, the protein production from linear mRNA decreased rapidly (with a >80% reduction on day 2), whereas the translation from circRNAs lasted 5-7 days (Fig. 3B). The prolonged protein translation is also consistent with our previous results using back-spliced circRNA reporter ^3^. It is also worth noting that, in this experiment, the protein production from circRNAs may also be suppressed because the cells reached the stationary phase at the end (Fig. 2B), as we only replaced the culture medium without splitting the cells during the entire week of the culture.

**Fig. 3.**
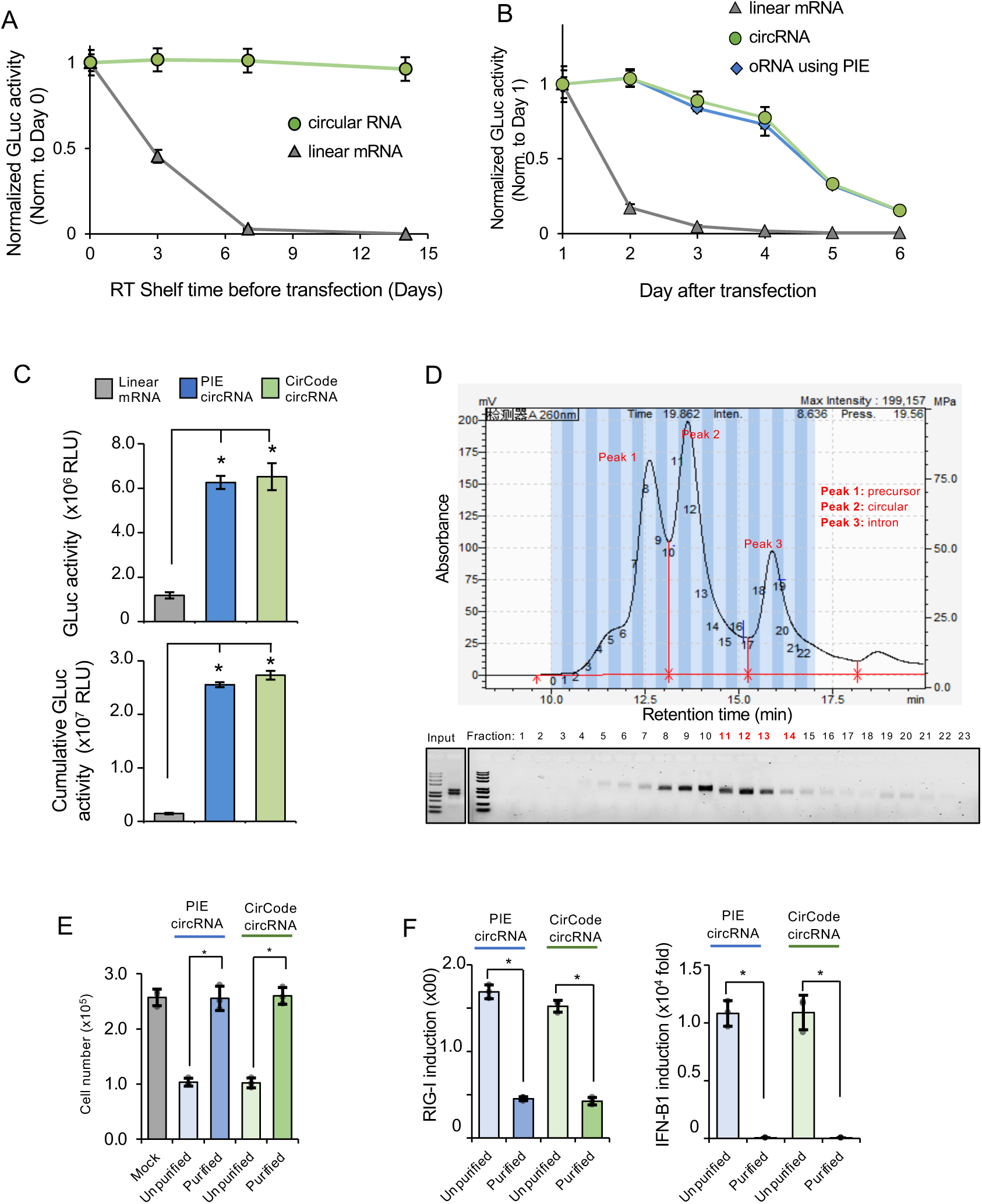
Stability and immunogenicity of circRNAs. (A) Thermostability of circRNA. Purified unmodified CVB3-Gluc circRNA and Gluc mRNA were diluted to the final concentration 100 ng/ul by nuclease-free water, and were stored at room temperature (25 ℃). RNA samples were collected at day 0, 3, 7 and 14, and stored at -80℃ for further analysis. All collected RNA samples were then transfected into 293T cells (100ng circRNA and mRNA were used per well in 96-well plate), and the luciferase activity in the media were measured at 24 hours after transfection. (B) Prolonged protein expression of circRNA in cells. Unmodified Gluc mRNA and CVB3-Gluc circRNAs generated through group II intron or PIE method were transfected into 293T cells (equimolar quantities of each RNA, equivalent to between 50 and 100 ng dependent on size, were used per well in 96-well plate). The cell culture media were fully collected and replaced at 1, 2, 3, 4, 5, and 6 days after transfection, and the luciferase activity in media were measured. (C) High protein expression of circRNA containing the CVB3 IRES and Gluc ORF. The luciferase activity in medium of 293T cells 24 h after transfection with CVB3-GLuc circRNA or unmodified GLuc mRNA (Top panel). Relative cumulative luciferase activity produced over 6 days by 293T cells transfected with CVB3-GLuc circRNA or unmodified GLuc mRNA (Bottom panel). (D) HPLC purification of CVB3-Gluc circRNA from spin column purified sample after IVT. The top panel is the HPLC chromatogram indicating the peak of precursor, circular and intron RNA, respectively. The bottom panel demonstrates the agarose gel of input and collected fractions. (E) Cell viability of A549 cells transfected with unpurified or purified CVB3 Gluc circRNAs generated by group II intron or PIE method. The cell viability was mesure 2 days after transfection. (F) The induced RIG-I and IFN-b1 gene expression of A549 cells at 24 h after transfection with unpurified or purified CVB3 Gluc circRNAs generated by group II intron or PIE method.

We further compared the protein production from the linear mRNAs and the circRNAs produced using the PIE method or the new CirCode system. As a control, the unmodified linear mRNAs were generated using IVT with the same coding sequences of Gluc, and transfected into 293 T cells in parallel with the two types of circRNAs. We found a more robust expression of proteins from both circRNAs compared to the linear mRNA (Fig. 3C, top), supporting the previous reports using PIE circRNAs ^9^. An even larger increase in protein production was observed in circRNAs when we measured the accumulated luciferase activity over the span of 6 days (Fig. 3C, bottom), presumably because of the superior stability of circRNAs. In addition, our data suggested that the circRNAs generated from two different methods showed similar ability in the direct protein translation.

### Purified circRNAs can direct robust translation of target proteins

The mRNA purity was found to be a key factor for the protein production and induction of innate immunity, as the removal of dsRNA by HPLC can eliminate immune activation and improves translation of linear nucleoside-modified mRNA ^20^. However, there are some debates on the immunogenicity of circRNAs. While an early report suggested that *in vitro* synthesized circRNAs are more prone to induce cellular immune response than the linear RNA ^21^, it was later reported that purification of circRNAs from byproducts of IVT and circularization reactions, including dsRNA, linear RNA fragments and triphosphate-RNAs, can eliminate the cellular toxicity and immunogenicity of the circRNAs ^4,5^. A recent study also suggested that the sequence identity and structure are the main determinant of cellular immunity of circRNAs, as the circRNAs produced by different methods showed different immunogenicity ^11^. To examine if the circRNAs produced using CirCode platform can induce innate immune response and cell toxicity, we purified the circRNAs with gel purification or HPLC (Fig. 3D), and measured if the circRNAs can induce cellular immune response upon transfection of circRNAs. We found that transfection of the unpurified circRNAs cause a significant amount of cell death, whereas the purified circRNAs did not show detectable cell toxicity compared to mock transfection (Fig. 3E). In addition, compared to the unpurified circRNAs that stimulated innate immune response by inducing RIG-I and IFN-B1 (interferon-β1), the purified circRNAs showed minimal immunogenicity (Fig. 3F), supporting the previous observation that the immunogenicity of circRNAs was mainly caused by the by-products from IVT reaction ^4,5^.

### LNP encapsulated circRNAs direct robust protein production in mouse

An important question for the therapeutic application of circRNAs is whether the circRNA production can be scaled up reliably and how is the reproducibility between different batches of production. Because the CirCode platform uses self-splicing intron for RNA circularization without the involvement of RNA ligase or additional co-activators, the procedure is relatively simple to scale up. To test the scalability of this system, we proportionally expanded the IVT and circularization reaction for 50 fold (from 20 μl to 1 ml), and found that the high circularization efficiency (∼70%) stayed essentially unchanged while the total amount of RNA products reached 7.5 mg in a single reaction (Fig. 4A). In addition, we found that the liquid chromatography production and purification of the circRNAs were scalable and highly reproducible across different batches (Fig. S3 and Fig. 4B), laying the ground for the *in vivo* application of the circRNAs.

**Fig. 4.**
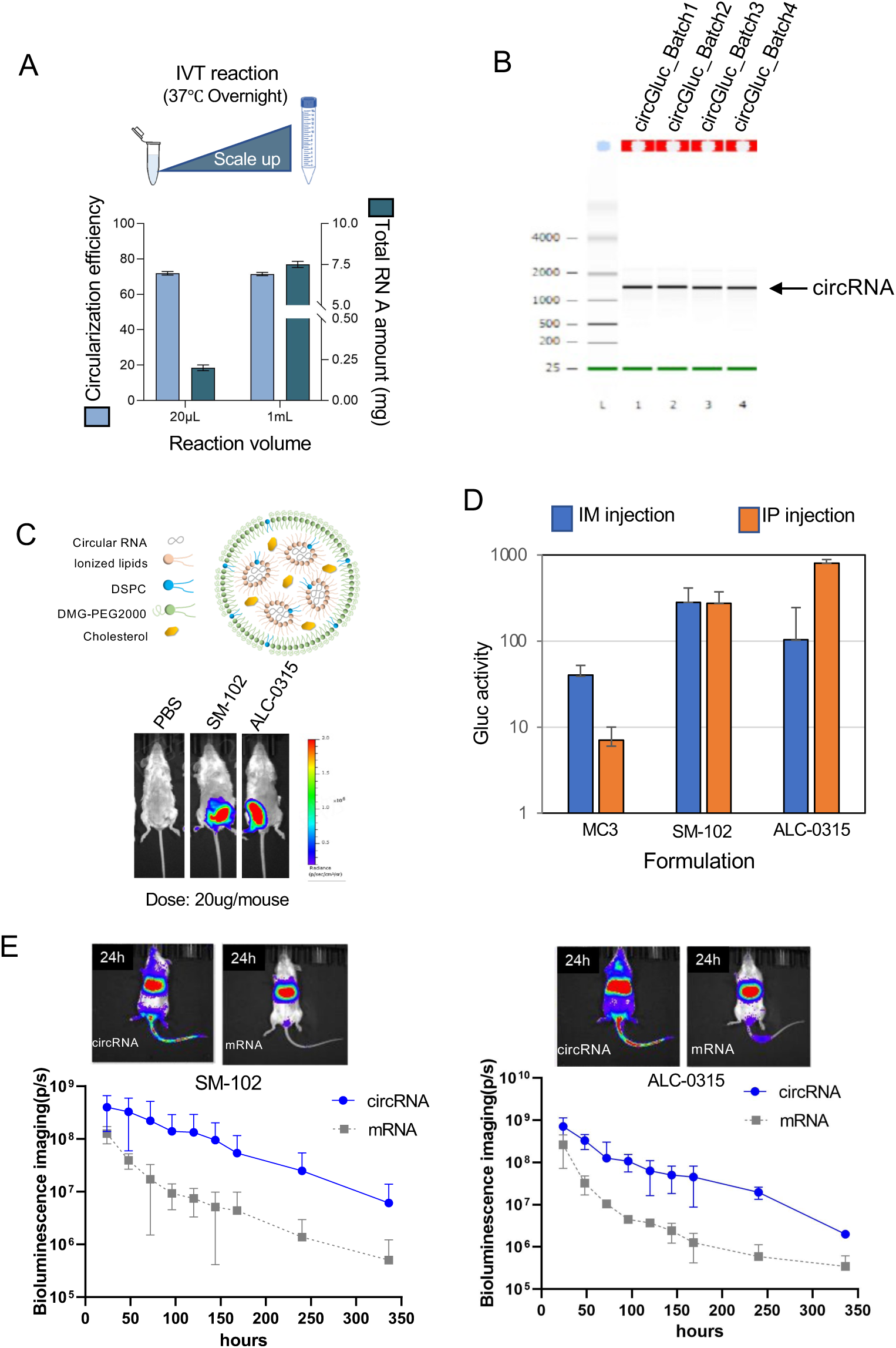
*In vivo* delivery of circRNA for protein expression in mice. (A) Scale up the production of circRNAs from 0.25 mg to ∼7.5 mg. (B) Reproducibility of the circRNA production. Four batches of Gluc circRNA purified from HPLC were analyzed using capillary electrophoresis with Agilent 2100 Bioanalyzer. (C) Schematic illustration of circRNA-LNP complex and particle size of CircRNA^Gluc^-LNP, and representative IVIS images of BALB/c mice administrated with 20ug CircRNA^Gluc^-LNP with two formulations by the intramuscular (i.m.) routes. Relative luminescence plot is shown and the scale of luminescence is indicated. (D) Gluc activity assayed from mice serum 24 hours post-injection of CircRNA^Gluc^-LNP with different formulation. (E) The unmodified circRNA or linear mRNA fully modified with N1-methylpseudouridine were packaged with two different LNP formulations and intravenously injected into mouse at 10 ug/mouse dosage. Both circRNA and linear mRNA have same ORF encoding encoding firefly luciferase (Fluc), and the protein production was followed as luciferase activity (bioluminescence) for 2 weeks. The representative images at 24 hours post injection were also showed as example.

We further generated circRNAs encapsulated with lipid nanoparticles (LNP) for their *in vivo* delivery (Fig. 4C). The circRNA in the aqueous solution was packed by ionizable cationic lipids, which formed a nanoparticle with other lipid components such as DMG-PEG2000 and cholesterol (see methods), resulting in a ∼95% encapsulate efficiency with the effective diameter at ∼80 nm. We have tested three different ionizable cationic lipids in our formulation to encapsulate the circRNAs encoding Gluc, and the resulting LNP-circRNAs were injected into BALB/c mice through intramuscular (IM) or intraperitoneal (IP) injection (n=3 for each experimental group). Three formulations using different ionizable cationic lipids (MC3, SM-102 and ALC-0315) were tested in this experiment. The expression of luciferase was assayed by the bioluminescence imaging of the animals (Fig. 4C) and the measurement of luciferase activity in serum (Fig. 4D). We found a robust expression of luciferase from two formulations of LNP-circRNAs using SM-102 or ALC-0315 as ionizable lipids, suggesting that the circRNAs produced through CirCode system can reliably induce *in vivo* protein expression.

To further assess the potential of using circRNAs as an alternative for mRNA therapy, we compared the efficiency for the *in vivo* protein expression from synthesized circRNAs and linear mRNAs. The unmodified mRNA is known to induce innate immunity that inhibit the protein translation and reduce the efficiency for therapeutic protein expression, which can be reduced by introducing base modifications ^22,23^. The N1-methylpseudouridine (m1Ψ) base modification, used in SARS-CoV-2mRNA vaccines, is the most effective modification in promoting the translation efficiency of synthesized mRNA ^24^. Therefore we compared the CirCode circRNA and fully modified linear mRNA with the same ORF sequence encoding firefly luciferase (Fluc). Using intravenous tail injection, the LNP-circRNA or LNP-mRNA were delivered systematically into mice, and the protein expression was followed with bioluminescence for 2 weeks. We found that the optimized circRNAs can provide a stronger and longer expression compared to modified linear mRNAs (Fig. 4E),

### Generation of a potential circRNA vaccine for SARS-CoV-2

We further tested the application of circRNAs in mRNA therapy by engineering the circRNAs encoding the receptor binding domain (RBD) from the S protein of SARS-CoV-2, which can potentially be used to produce mRNA vaccine. Based on previous reports, two different antigen designs were constructed into the circRNAs (Fig. 5A top) ^25^. The first one used a single RBD fused with the foldon from T4 fibritin that enables trimerization of RBD ^26,27^. The other used a RBD dimmer that was shown to induce strong antibody production in pilot study ^25,28^. We have designed the coding sequences of both proteins with an engineered IRES, and constructed the circRNA vectors. The purified circRNAs were validated with capillary electrophoresis (Fig. 5A, left), and translation of the circRNA-encoded proteins was further validated by transfecting into 293 cells and detected by western blot antigen production (Fig. 5A, right).

**Fig. 5.**
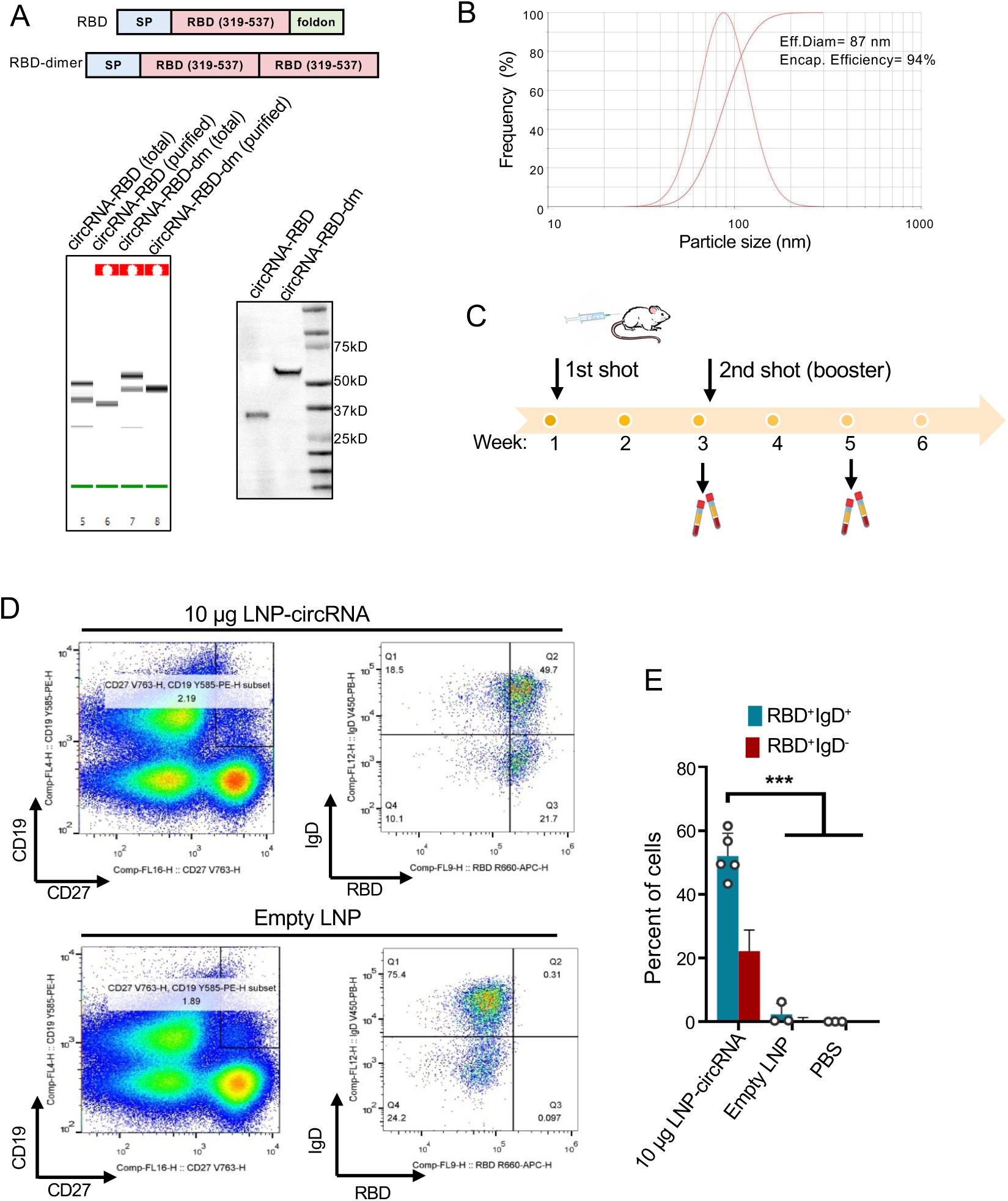
Design circRNAs for Covid-19 vaccine. (A) circRNA RBD and circRNA RBD dimer purified from HPLC were analyzed using capillary electrophoresis with Agilent 2100 Bioanalyzer (left bottom), the protein expression of cell culture medium from circRNA-RBD or circRNA-RBD dimer was determined by western blot at 24hr after transfection. (B) The particle size and encapsulate efficiency of circRNA RBD-LNP complexes. (C) Schematic diagram of the circRNA RBD-LNP vaccination process in BALB/c mice and serum collection schedule for antibody analysis (D) Represented unvaccinated and vaccinated cohorts are shown for RBD specific B cell responses. (E) FACS analysis results showing the percentages of RBD specific B cell.

We used two different formulations to produce the LNP-circRNA particles, and achieved high encapsulation efficiency (>90%) with typical nanoparticle size at 90-100 nm (Fig. 5B showing the representative result using SM-102 formulation). The LNP-circRNAs were inoculated into the BALB/c mice with two separated IM injections, and the blood samples were collected at 2 weeks after each inoculation for further analysis (Fig. 5C). We tested several formulations and circRNA designs, however the results from the LNP-circRNA-RBP using SM-102 formulation were further examined for a better comparison with published results. A flow cytometry antibody panel was designed to identify naïve B cells (CD19+/IgD+/CD27−), total memory B cells (CD19+/CD27+), including an unswitched IgD+ population and a switched IgD− population, plasma cells (CD19+/IgD−/CD38+/CD27+), and transitional B cell (CD19+/IgDdim/CD38+). To determine whether LNP-circRNA-RBP vaccination induced the activation and expansion of antigen-specific B cells, we measured the frequency of RBD-binding B cells using Alexa 647 labeled RBD (RBD-Alexa 647). A large fraction of RBD-specific lymphocytes were detected in the memory B-cells (CD19+/CD27+), including an RBD specific switched B-cell population (CD19^+^CD27^+^ IgD^−^ RBD^+^) and an RBD specific unswitched memory B-cell population (CD19^+^CD27^+^ IgD^+^ RBD^+^) (Fig. 5D, 5E), indicative of a long-lasting immune response induced by antigen. In particular, >70% of CD19^+^CD27^+^ B-cells expressed RBD-specific antibody (i.e., RBD^+^), and ∼50% of the CD19^+^CD27^+^ B-cells were also IgD+/RBD+ (Fig. 5E), suggesting a strong RBD-specific memory B cell response elicited by the LNP-circRNA vaccination. As a control, the injection of LNP alone did not induce activation of RBD-specific memory B cells, confirming that the observed response was specifically induced by the circRNA.

To test the activity of the RBD antibody in mouse serum, we next performed an antibody blocking assay to examine if the mouse serum can block the binding of Alexa 647-labeled RDB to the 293 T cells stably expressing human angiotensin I-converting enzyme 2 (hACE2) (Fig. 6A). We found that the serum can effectively block the binding of three different RBD variants (RBD_wild type_, RBD_delta_, and RBD_omicron_) to the cell surface at a modest dilution (1:20). Given that we used a pretty high concentration of RBD (0.5 μg/ml), this protection is quite impressive. Even in a very high dilution ratio (1:200), the serum still completely blocked the binding of the wild-type RBD and significantly reduced the binding of the RBD of the Delta variant. However, the serum did not block the binding of the omicron RBD at the same dilution ratio, suggesting a diminished blocking activity for the omicron variant. This result is consistent with the finding that the current mRNA vaccines based on the wild type S protein are weak in protecting against the omicron variant ^29^.

**Fig. 6.**
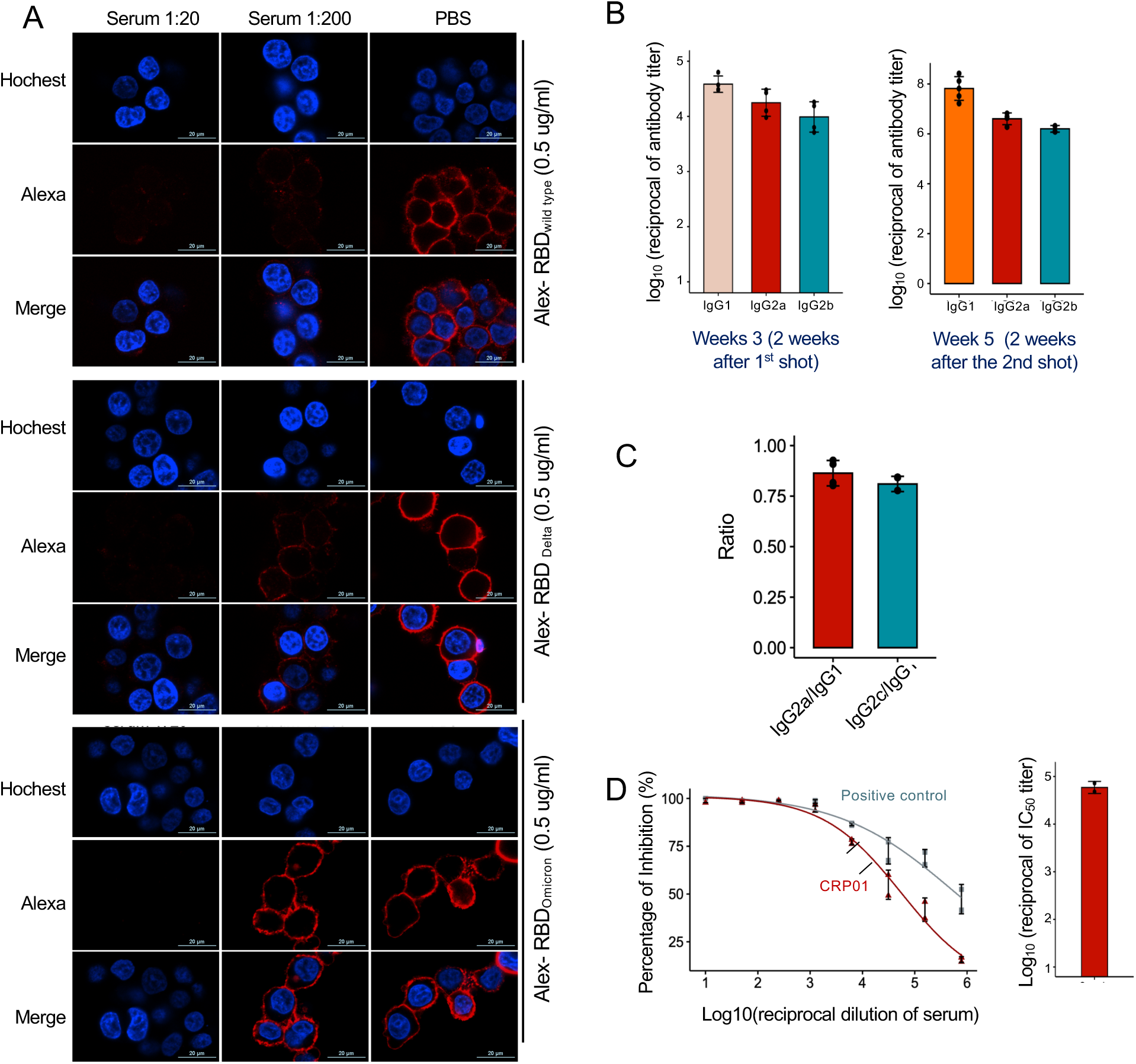
Analysis of neutralizing antibody generated by circRNA vaccine. (A) The competitive binding of RBD and immunized mouse serum to 293T cells expressing hACE2 *in vitro*. Alex-RBD: Alexa 647 labeled RBD. (B) Measurement of RBD-wt specific IgG1, IgG2a and IgG2c endpoint titers in mice at 2 weeks after 1^st^ shot and 2 weeks after 2^nd^ shot. (C) Measurement of RBD-wt specific IgG2a/IgG1 and IgG2c/IgG1 ratios. (D) Neutralization assay of lentivirus-based SARS-CoV-2 pseudovirus with the sera from mice immunized with 10ug circRNA RBD-LNP.

### circRNA mRNA as vaccine for SARS-CoV-2

We further measured the antibody titers after each inoculation, and found a robust production of IgG production against RBD (Fig. 6B-6C). The binding antibody titer was close to 10^5^ at two weeks after the first shot and reached near 10^8^ after the second shot, which is ∼10 times higher than the reported mouse results using the LNP encapsulated linear mRNAs with a similar formulation ^30^ or using another circRNA design ^31^. We further conducted a detailed comparison with the linear mRNAs

In addition, the ratio of IgG2/IgG1 is around 0.75, suggesting a balanced Th1/Th2 immune response. Similar immune response was previously reported using linear mRNA vaccine mRNA-1273 ^30^, implying circRNA vaccine may have a low risk in vaccine associated enhanced disease and a potential good safety profile. Finally, we measured the pseudo-neutralizing antibody titer using the protection assay for SARS-CoV-2 pseudovirus, and found a strong protection with a pseudo-neutralizing titer close to 10^5^ (Fig. 6D). Collectively, our preliminary data in the mouse model suggested that the CirCode platform had a strong performance in the design and generation of circRNA vaccines against SARS-CoV-2, which can potentially be expanded to the vaccine development for other viral pathogens.

## Discussion

The approval of mRNA vaccines against Covid-19 pandemics has pushed the mRNA therapy into the front stage. In addition to the vaccine, mRNA drugs had shown potential in the treatment of a variety of diseases, including cancers, rare genetic disorders, and neuronal diseases ^1^. However, linear mRNAs are intrinsically unstable, which limited their application. Several features of circRNA make them an appealing candidate as the new generation of mRNA drugs. First, circRNAs have increased stability both *in vitro* and *in vivo*, which enables a prolonged expression of target genes. Second, the chemical modification of circRNAs seems to be unnecessary, as the unmodified circRNAs can be reliably translated without inducing the innate immune response (Fig. 3E) ^4,5^. In addition, circRNAs use IRESs or IRES-like sequences to initiate translation in a cap-independent fashion ^3,19^, which may allow cell-type specific translation through cellular *trans*-acting protein factors ^4,19,32^. Despite these advantages, there are technical challenges to overcome before circRNAs become a general gene expression platform, including circRNA manufacturing, purification and quality control, optimization of IRES mediated translation, as well as *in vivo* delivery. We have developed a new general platform for circRNA design and production, which can be integrated with the sequence design and IRES optimization to achieve a high efficiency of protein translation from circRNAs.

The inverse splicing of both group I and group II self-catalyzed introns have been discovered more than two decades ago to produce a small amount of circRNAs ^8,10,33^. However, the circularization efficiency is very low in these early studies, and the circRNAs can only be produced from their intrinsic exon sequences. Recently the PIE method has been optimized to produce circRNAs containing customized target sequences ^9^, however a large fragment of the exons adjacent to the group I intron have to be included in the final circRNA product as a scar sequence. In this study, by screening a series of group II introns and analyzing the intron structures, we were able to engineer a new split-intron system to produce circRNAs containing a short foreign fragment of 12 nt or do not contain any foreign fragment (i.e., scarless). The main reason for the scarless inverse self-splicing is that we were able to rearrange the structure of group II intron in a modular fashion to make it accommodate different exon sequences at the splice junction. In addition, the target sequence seems to play a less role in affecting the folding of group II intron, because our method can achieve decent circularization efficiency (>50%) for most genes. With additional optimization, we can routinely obtain a circularization efficiency of 60-90% with different designs of the junction sequences.

Because of superb stability and low immunogenicity, the base modification seems to be unnecessary for circRNAs. In addition, circRNA production does not require the step of 5’ capping, which can significantly reduce the complexity and cost for circRNA production. The new method in this study has enabled co-transcriptional circularization without the supplement of extra GTP, which further streamlines the production process.

The emerging new variants of SASR-coV-2 presented a challenge for new mRNA vaccines. Because the levels of antibodies induced by mRNA were found to decrease gradually, the repeat booster of vaccination is usually needed for effective protection ^34^. Using circRNA as a new vaccination platform has been shown to be a promising alternative to the linear mRNA vaccine ^31^, probably because of its superb stability. Compared to the linear mRNA, the antigen production from cap-independent translation of circRNA may peak slower but last longer, resulting in more antigen production by accumulative protein expression. It remains unclear how this different dynamic of antigen expression can affect the antibody response, however our early results showed a very strong induction of memory B-cells. Additional experiments will be needed to optimize the circRNAs for vaccine production, including the comparison of different antigen designs and optimization for the formulation.

There are still many technical questions remain to be solved for circRNAs as a new therapeutic reagent, including the sequence optimization for circRNA translation and the specific methods for *in vivo* delivery. Although the canonical IRESs that drive circRNA translation is mainly derived from viral sequences, recent studies found that many additional sequence elements can function as IRES-like elements or regulatory elements to promote IRES activity ^19,35^. Therefore the sequence optimization for circRNA translation will likely be different from the optimization of UTR sequences in linear mRNAs. In addition, the superior stability of circRNAs may make them more tolerable to additional formulation methods of RNA delivery. We found that most formulations for mRNA delivery can efficiently deliver circRNAs, however it is both intriguing and important to study if certain methods unfit for linear mRNA delivery may actually work well for circRNAs. Overcoming these technical hurdles will certainly improve our understanding on the translation regulation of circRNAs, as well as the circRNA transport between or inside cells.

In summary, we have developed a new system for efficient and scalable production of circRNAs in a large scale, which is suitable for industry level applications. The resulting circRNAs can be engineered to direct robust protein translation, providing a new platform of mRNA therapy with improved stability and reduced immunogenicity. In addition, this system can also be used to generate various circRNAs with different therapeutic activities, such as miRNA sponge or RNA aptamers. The continuous improvement of this platform should help to take circRNA technology into various clinic applications in the near future.

## Methods

### Plasmid construction

The fragments of the group II intron in *Clostridium tetani* (CTE) and IRES sequences were chemically synthesized from GENEWIZ, and selected mutations were introduced in the synthesis to make it recognize different exons. Different protein coding fragments (ORF) were amplified by PCR and merged with the IRES fragments using PCR reactions. These fragments were cloned into the NheI and XbaI digested backbone containing T7 RNA polymerase promoter and terminator using Gibson assembly (see supplementary data 1 for the sequences of these DNA segments and primers).

### RNA synthesis and circularization

The plasmid DNAs were linearized with XbaI digestion and purified with Post PCR cleanup kits from Qiagen. The linearized DNAs were used as a template for *in vitro* transcription with T7 RiboMAX™ Large-Scale RNA Production System (Promega, P1320) in the presence of unmodified NTPs. After DNase I treatment, the RNA products were column purified with RNA Clean and Concentrator Kit (ZYMO research, R1013) to remove excess NTP and other salts in IVT buffer, as well as the possible small RNA fragments generated during IVT. In some experiments, the purified RNA was further circularized in a new circularization buffer. The RNA was first heated to 75°C for 5 min and quickly cooled down to 45°C, after which a buffer including indicated magnesium and sodium was added to a final concentration: 50mM Tris-HCl at pH 7.5, 50/100mM NaCl, 0-40mM MgCl_2_, and was then heated at 53°C for indicated time for circularization. The best-optimized reaction condition including concentration of magnesium and sodium, and incubation time at 53°C, was selected for further experiments.

### CircRNA identification

For the poly A tailing and RNase R treatment, the total RNAs from IVT were purified by RNA cleanup columns, and then treated with *E. coli* Poly A Polymerase (NEB, M0276S) following the manufactory instruction. This step will add a poly-A tail to the free end of the unspliced linear RNA precursor. After Poly A-tailing, the purified RNAs were digested by RNase R exoribonuclease (Lucigen, RNR07520) following the manufacturer’s instructions, and enriched circRNAs were purified by column.

For the RNase H nicking assay, the RNase R enriched circRNAs were incubated with a 24-nt ssDNA probe at 1:20 ratio, The RNase H buffer was added to the DNA-RNA mixture immediately. Subsequently, the mixture slowly cool to room temperature. After annealing, RNase H (Thermo Scientific, EN0201) was added to the mixture for 20min at 37 °C. The sequence of the ssDNA probe is 5’-TGGTGCTCGTAGGAGTAGTGAAAG-3’.

For all the gel electrophoresis, the total RNA from IVT samples was separated on a low melting point agarose gel (Sigma Aldrich, A4018) at 120V using ice-cold DEPC-treated MOPS butter. To gel purify the circRNA, the circRNA band was cut and purified with Zymoclean Gel RNA Recovery Kit (ZYMO research, R1011).

To analyze the circRNA with sequencing, the gel-purified RNA was reverse transcribed into cDNA using a PrimeScript RT Reagent Kit with random primers (TAKARA, RR037B), followed by PCR with primers that can amplify transcripts across the splice junction. The PCR products were sequenced using Sanger sequencing to validate the backsplice junction of the circular RNA.

### HPLC purification and electrophoresis of RNA

To obtain the high-quality circRNA, spin column purified DNase I-treated RNA from IVT was resolved with high performance liquid chromatography (HPLC). For a small-scale preparation with SHIMADZU LC-20A (Kyoto, Japan), 40μg RNA was loaded onto a 4.6×300 mm size exclusion column (Waters XBridge, BEH450A, 450Å pore diameter, 3.5μm particle size) and eluted with mobile phase containing 10mM Tris, 1mM EDTA, 75mM PB, pH7.4 at 25°C with flow rate 0.5ml/min. For a large-scale preparation with Sepure SDL-30 (Suzhou, China), 4mg RNA was loaded each run onto a 30×300mm SEC column (Sepax, SRT SEC-1000A, 1000Å pore diameter, 5.0μm particle size, Suzhou, China) with mobile phase containing 10mM Tris, 1mM EDTA, 75mM PB, pH7.4 at 25°C with flow rate 10ml/min. Fractions were collected as indicated and testified with agarose gel electrophoresis.

Circular RNAs purified from large-scale production were further analyzed with capillary electrophoresis with Agilent 2100 Bioanalyzer in the RNA mode. Samples were diluted to a propriate concentration and analyzed according to the manufactory’s instructions.

### Measurement of the translation products from circRNAs

Cells were seeded into 24-well plates one day before transfection. The purified circRNAs are transfected into cells using Lipofectamine Messenger Max (Invitrogen, LMRNA001) according to the manufacturer’s manual. After transfection, cells were cultured at 37°C for 24h. The cell lysis and supernatant were collected for luminescence assay using the Dual-Luciferase® Reporter Assay System (Promega, E1910).

### LNP Production Process

The circular RNA was encapsulated in a lipid nanoparticle via the NanoAssemblr Ignite system as previously described ^30,36^. In brief, an aqueous solution of circRNA at pH 4.0 is rapidly mixed with a lipid mixture dissolved in ethanol, which contains different ionizable cationic lipid, distearoylphosphatidylcholine (DSPC), DMG-PEG2000, and cholesterol. The ratios for the lipid mixture are MC3:DSPC:Cholesterol:PEG-2000=50:10:38.5:1.5 for formulation 1, SM-102:DSPC:Cholesterol:PEG-2000 = 50:10:38.5:1.5 for formulation 2, and ALC-0315:DSPC:Cholesterol:ALC-0159 = 46.3:9.4:42.7:1.6 for formulation 3. The resulting LNP mixture was then dialyzed against PBS and stored at -80 °C at a concentration of 0.5 μg/μl for further application.

### Administration of LNP-circRNAs in mice

Female BALB/C mice aged 8 weeks were purchased from Shanghai Model Organisms Center. 20 μg of LNP-circRNAs in PBS were administrated into mice intramuscularly with 3/10 insulin syringes (BD biosciences). The serum was collected 24 hours after the administration of LNP, and 50 μl serum was used for Luciferase activity assay *in vitro*. Bioluminescence imaging was performed with an IVIS Spectrum (Roper Scientific). 24 hours after Gluc-LNP injection, 2 mg/kg of Coelenterazine (MedChemExpress,MCE) was administrated to mice intraperitoneally. Mice were then anesthetized after receiving the substrates in a chamber with 2.5% isoflurane (RWD Life Science Co.) and placed on the imaging platform while being maintained on 2% isoflurane via a nose cone. Mice were imaged 5 minutes post substrate injection with 30 seconds exposure time to ensure the signal was effectively and sufficiently acquired.

For immunogenicity studies, 8-week-old female BALB/c mice (Shanghai Model Organisms Center, lnc) were used. 10μg of CircRNA-RBD-LNP were diluted in 50μl 1XPBS and intramuscularly administrated into the mice’s same hind leg for both prime and boost shots. Mice in the control groups received PBS and empty LNPs. The blood samples were collected 2 weeks after prime and boost shots. Mice spleen were also harvested at the endpoint (2 weeks after boost) for immunostaining and flow cytometry.

### B cell surface staining for flow cytometry

The spleenocyte suspensions from whole spleens were generated using a tissue dissociator (RWD Life Science) followed by 70-μm filtration. Cells were isolated and resuspended in R10 media (RPMI 1640 from Gbico, supplemented with Pen-Strep antibiotic, 10% HI-FBS, 1%Glutamax, and1% HEPES) followed by density gradient centrifugation using Fico/Lite-LM medium (R&D Systems). Cells were washed and stained with Fixable Viability Stain 510 (BD Pharmingen) at 4℃ in the dark at least 5 min in brilliant stain buffer. Cells were then washed with FC buffer (PBS supplemented with 2% HI-FBS and 0.05% NaN3) and resuspended in Fixation Buffer (BD Pharmingen)for 5 min at 4℃ in the dark. Cells were then blocked with Fc Block (BD, clone 2.4G2; 1:100) at 4℃ for 5 min, cells were labeled with the following antibodies: CD19-PE (BD), CD38-BV421(BD), CD27-BV786(BD), IgD-BV650(Biolegend), RBD-Alexa 647 in brilliant stain buffer (BD) for 20min. Cells were then washed and resuspended in FC buffer before running on a LSRFortessa flow cytometer (BD).and anlayzed using FlowJo software version 10.

### ELISA

Elisa plates (Corning) were coated with 100ng per well of recombinant Spike RBD protein in 1×coating buffer (Biolegend) overnight at 4°C. After standard washes and blocks, plates were incubated with serial dilutions of heat-inactivated sera for 2 hours at room temperature followed by standard washes. Anti-mouse IgG1, IgG2a, or IgG2c-horseradish peroxidase conjugates (Abcam) were used as secondary antibodies. After 1 hour incubation, plates were washed and 3,5,3′5′-tetramethylbenzidine (TMB) substrate (Beyotime) were used as the substrate to detect the antibody specific signal. The reaction was stopped using stop solution (Beyotime) and the final result was read at 450 nm on an Elisa plate reader (Biotek). End-point titers were calculated as the dilution that emitted an optical density exceeding 4 times of background and calculated in Graphpad Prism software version 8 and R-Studio.

### Lentivirus-based pseudovirus-neutralization assay

SARS-CoV-2 lentivirus-based pseudovirus encoding a luciferase reporter (Genwiz) was used for neutralization assay following a previously described method (cite). In brief, serial dilutions of heat-inactivated sera were mixed with pseudoviruses, incubated, and then added to hACE-2 and human transmembrane protease serine 2 (TMPRSS2) co-expressing HEK293 cells followed by a period of incubation time. Cells in each well were lysed and measured for luciferase activity (in relative light units (RLU)). Neutralization was calculated considering uninfected cells as 100% neutralization and cells infected with only pseudovirus as 0% neutralization. IC50 titers were determined using a log (agonist) vs normalized-response (variable slope) nonlinear function in Graphpad Prism software version 8 (GraphPad) and R-Studio.

### Confocal Immunofluorescent analysis of RBD binding

Dilutions of heat-inactivated sera were mixed with Alexa-647 labeled RBD protein followed by 5 minutes at room temperature. hACE-2-expressing HEK293 cells were seeded on a Nunc Lab-Tek Chamber Slide system (Thermo Fisher) and cultured overnight. Cells were washed and stained with Hoechst for nuclear staining followed by staining with the mixture of sera and Alexa-647 labeled RBD. Cells were washed and imaged using confocal microscopy (Sony).

## ACKNOWLEDGEMENT

The authors want to thank Dr. Leaf Huang for his suggestions and comments on the manuscript. This work is partially supported by the National Natural Science Foundation of China to Z.W. (91940303 and 31730110) and Y.Y. (31870814). Z.W. is also sponsored by the type A CAS Pioneer 100-Talent program, Y.Y. is also sponsored by the Youth Innovation Promotion Association CAS, SA-SIBA Scholarship Program and Shanghai Science and Technology Committee Rising-Star Program (19QA1410500). Z.W. is also supported by the Starry Night Science Fund at Shanghai Institute for Advanced Study of Zhejiang University (SN-ZJU-SIAS-009).

---

**Fig. S1.**
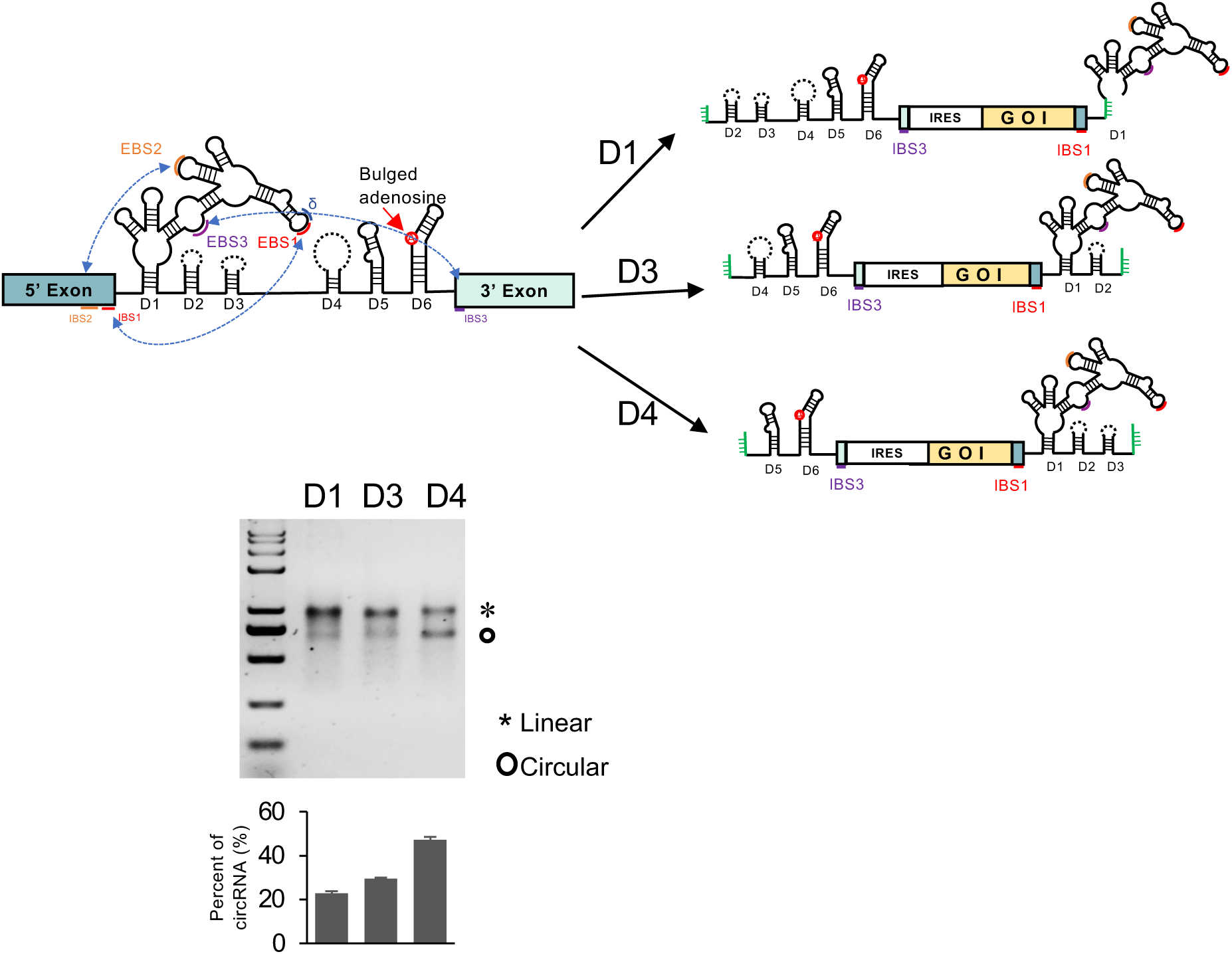
Analysis of circularization efficiency of Group II intron with different split sites. Top, schematic diagram for the design of CirCode system with different split sites. The autocatalytic self-splicing group II intron was split into two fragments at the D1, D3, or D4 domain, and a customized exons containing IRES and coding region of a gene of interest (GOI) were inserted between the split introns. Bottom, domain, and a customized exons containing IRES and coding region of a gene of interest (GOI) were inserted between the split introns. Bottom, the circRNA is in vitro synthesized and analyzed with agarose gel.

**Fig. S2.**
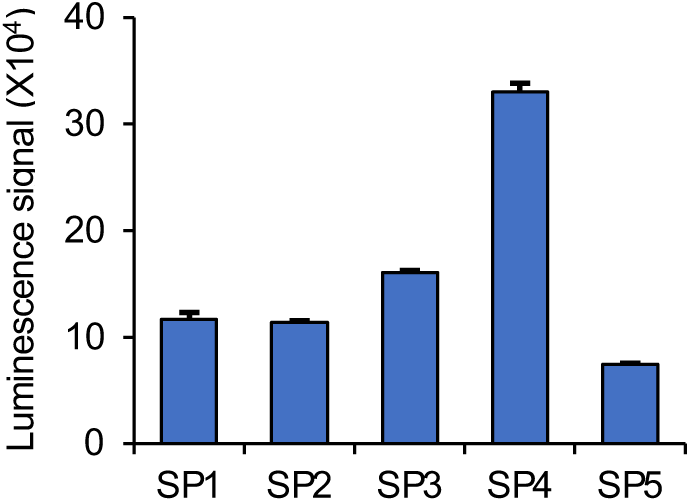
Analysis of protein expression from circRNA with different spacer regions. The protein expression from circRNA containing CVB3 IRES and Gluc ORF with different spacer regions (from Fig. 2B) at 48 hours after transfection.

**Fig. S3.**
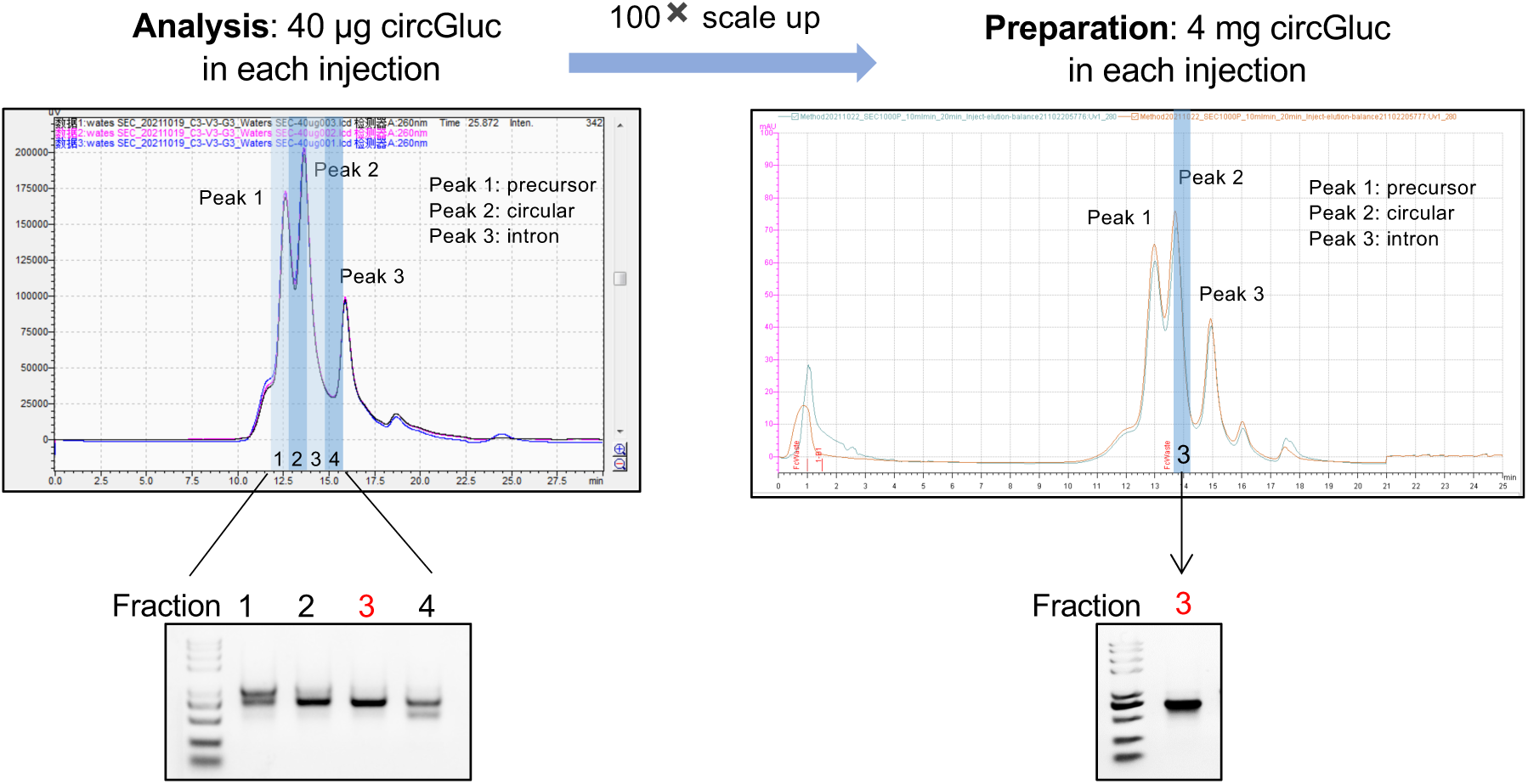
Analytical and scale-up SEC of circRNA from splicing reactions by HPLC. Left panel indicates the analytical HPLC chromatogram of 40μg circGluc with three injections (left top) and agarose gel of collected fractions (left bottom). Right panel demonstrates HPLC scale-up chromatogram of 4mg circGluc with two injections (right top) and agarose gel of collected fraction (right bottom)

